# Age-related differences in episodic memory precision are robust across time of day, chronotype, and testing format

**DOI:** 10.64898/2026.09.28.755202

**Authors:** Julia S. Maybury, Jon S. Simons

**Affiliations:** Department of Psychology, University of Cambridge, United Kingdom

## Abstract

Older adults typically exhibit reduced episodic memory precision, while maintaining relatively preserved retrieval success. However, little is known about whether these age-related differences are influenced by internal biological factors (e.g., chronotype) or external testing conditions (e.g., experiment format, time of day).

To address this question, younger and older adults completed an object-location memory task online, either in the morning or evening, and either with or without video-based supervision. Memory performance was assessed using mean absolute error and Bayesian hierarchical mixture modelling, which dissociated memory precision (κ₁) from the probability of memory-based responding (θ₁).

Immediate memory performance was high across age groups, irrespective of time of day. Greater encoding error predicted greater retrieval error, and participants tested in the video condition exhibited lower recall error and higher recognition performance than those tested without supervision. Consistent with prior studies, older adults exhibited reduced memory precision relative to younger adults, whereas retrieval success remained largely intact. However, Bayesian Hierarchical Modelling provided no evidence that time of day, experiment format, or their interactions with age influenced either memory precision or retrieval probability. Older adults reported stronger morning preferences than younger adults, and participants generally selected testing times that aligned with their chronotype. Nevertheless, alignment between chronotype and testing time did not modulate memory performance. The results indicate that age-related differences in episodic memory precision are robust across variations in testing context, including time of day, experiment format, and individual differences in chronotype.

More broadly, the results support the feasibility of remote and unsupervised testing for memory precision research and suggest that chronotype and testing time are unlikely to be major sources of variability in immediate object-location memory performance.

## Introduction

An extensive literature has demonstrated significant age-related deficits in retrieving episodic memories, recollections of personal events tied to a specific time and place [1,2].

Although much has been learned, continuous memory paradigms offer an opportunity to further our understanding of age-related changes in episodic memory [3–5]. In such tasks, performance is assessed on a continuum, enabling researchers to dissociate behaviourally *memory accessibility*, a quantitative measure of performance (the probability of successfully retrieving a memory), from *memory precision*, a qualitative measure (the specific details of the retrieved memory representations).

Prior studies have implemented continuous object-feature recall tasks, which consist of a series of study-test phases separated by a variable delay period [6–8]. In the study phase of a long-term memory version of the task, for example, participants first encode stimulus displays consisting of everyday objects placed on a circular space and overlaid on a scene background.

During the test phase, participants are tested on perceptual features such as an object’s colour, orientation, or location using a continuous response dial with which they try to reproduce the object’s studied perceptual features as precisely as they can. Memory precision is assessed by the angular difference or error between the response and the target value. Probabilistic mixture models can be fit to the response error, enabling one to dissociate retrieval failure from imprecise retrieval of the target location.

While age-related reductions in memory precision are observed across different perceptual features (colour, orientation, location), the probability of successful recall from memory appears to be less influenced by age [7]. However, increasing task demand may reduce this dissociation. For example, when assessing the influence of different types of retrieved information, Korkki et al. [7] found retrieval success to be comparable between younger and older adults in the colour feature condition. In contrast, age-related decreases were observed in the orientation condition, which was the feature that younger adults found most challenging to retrieve successfully.

Performance on object location memory tasks may not only be influenced by age related differences in memory but also by contextual factors such as the interaction between time of day and chronotype, an individual’s sleep-wake cycle driven by the circadian system [9]. Chronotype can be measured using a Morningness-Eveningness Questionnaire (MEQ), a 19-item self-reported assessment of questions related to preferred wake/sleep times, timing of physical and mental activities and alertness throughout the day [10]. The synchrony effect refers to the phenomenon that individuals tend to perform better on tasks when they are completed at times of day that align with their natural circadian rhythms, or chronotype [11,12]. Notably, the effect is more pronounced in older adults, who exhibit a greater decline in performance during non-optimal times of day compared to younger adults [11,13]. Older adults often exhibit enhanced cognitive performance on tasks that require efficient attentional control in the morning, rather than in the afternoon or evening. In contrast, younger adults typically perform better in the afternoon or evening than in the morning [14]. Therefore, age differences in encoding at optimal and non-optimal times may influence the consolidation and retrieval of information [15,16].

Consequently, age differences in cognitive functioning throughout the day and the alignment between testing time and an individual chronotype are important contextual factors to examine and control.

In addition to when the task is performed, the effects on performance of where and how the task is administered may be influenced by age. Recent studies have demonstrated that unsupervised (non-video) online testing exhibited moderate reliability compared to in-person testing when older adults completed tasks measuring cognitive functions such as processing speed, executive functioning, and spatial working memory [17]. However, because performance on memory tasks is influenced by many factors such as attention, technology, and environmental distractions, careful consideration is required during experimental design to ensure the reliability of results. One way to potentially control these influences and still benefit from remote, online experimentation is video-based supervision. For example, using a Remote Guided Testing (RGT) protocol (a live video connection between researcher and participants) mitigated data quality issues, restricted external distractions, and ensured the integrity of cognitive metrics compared to unsupervised web-based testing in young adults [18]. An additional study demonstrated that core cognitive performance indices (such as error rates and accuracy) are consistent between unsupervised web testing and traditional supervised settings [19]. However, they noted subtle variances in reaction times and other performance measures that often occur due to hardware differences (e.g., varying monitor refresh rates, browser delays) and environmental distractions.

More age specific, Cyr et al. [20] explored how healthy older adults (aged 65+) handle the shift from supervised to unsupervised at home computerised cognitive task testing. Using a within-subjects design, they found accuracy and cognitive performance remained highly comparable across both settings, but supervised environments allow an experimenter to clarify complex instructions and manage technical anxiety. Just as they found no specific effects of the testing environment, so too they found no correlation of performance with computer familiarity or attitudes nor with paper and pencil equivalents. Although prior studies demonstrate consistent benefits to online testing, it remains unclear whether younger and older adults are equally robust to differences in supervised versus unsupervised testing, in particular for sensitive cognitive measures such as memory precision tasks. This is important to validate as online experiments provide an accessible, efficient, and affordable means of testing. One aim of the current study, therefore, is to measure high-precision memory tasks in both online unsupervised and supervised video conditions to examine how experiment format may affect performances across age groups, particularly in precision tasks which have repeatedly demonstrated age differences.

Accordingly, the study examined whether age-related differences in episodic memory precision were stable or context-dependent, examining both internal biological context (chronotype) and external context (e.g., experiment format, time of day). First, we explored the influence of internal biological factors by testing whether age differences in memory precision varied as a function of time of day (morning vs. evening) and whether performance was influenced by circadian alignment (synchrony effect). In addition, we examined whether these age-related differences were robust across external task contexts, comparing memory performance across two experiment formats (video vs non-video). Together, these internal and external factors evaluate whether age differences in memory precision are consistent across contexts or whether they could be shaped by biological and/or environmental conditions under which the memory is encoded and retrieved.

It is hypothesised, based on previous studies, that older adults will show reduced precision but intact retrieval success, independent of time of testing or the experiment format. However, consistent with the synchrony effect, older adults are expected to exhibit better memory performance in the morning relative to the evening, whereas younger adults are expected to show the opposite pattern, performing better in the evening. In addition, overall, participants are likely to exhibit lower memory performance in the non-video (unsupervised) compared to the video (supervised) experiment format, with greater deficit for older adults.

Examining chronotype, as measured by the MEQ, provides a theoretical explanation for the predicted time-of-day effects. Across both age groups, it is predicted that memory performance will be enhanced when testing time is in synchrony with participants’ chronotype. Specifically, individuals with more morning-type chronotypes (higher MEQ scores) are expected to perform better in the morning, whereas individuals with more evening-type chronotypes (lower MEQ scores) are expected to perform better in the evening. Therefore, across both age groups, it is predicted that greater similarity between an individual’s MEQ score and the time of testing will be associated with enhanced memory performance. This synchrony effect is expected to be more pronounced in older adults, given age-related shifts towards morningness and increased susceptibility to performing worse at off-peak times [11,13].

## Methods

### Participants

Participants were recruited online through Prolific between 24 April and 19 December 2024, and consisted of 123 younger-adults (aged 18-30 years) and 119 older-adults (60-80 years). The between-subject experiment was conducted online in either the morning or evening, using one of two formats: supervised (via video) and unsupervised (non-video) format. Seven participants from the video condition and three participants from the non-video condition were excluded from subsequent analyses as a result of either poor internet connection during the video session, poor recognition memory (hit rate > 3 SD below group mean), or learning or recall mean absolute error (>3 SD below the group mean). Therefore, for the video condition, the older adult sample analysed was *N* = 71 (age *range*= 60-78 years, *M =* 65.4, *SD =* 4.41) and the younger adult sample was *N* = 73 (age *range* = 18-30 years, *M =* 24.8, *SD =* 3.74). For the non-video condition, the older adult sample analysed was *N* = 42 (*range* = 60-80 years, *M =* 66, *SD =* 5.28) and the younger adult sample *N* = 46 (age *range*= 18-30 years, *M =* 26.2, *SD =* 3.40). All participants were United Kingdom citizens, fluent or native English speakers, with normal or corrected-to-normal vision. They also reported no current use of medications to treat symptoms of depression or anxiety, no known attentional problems (ADHD/ADD), and no diagnosis of dyslexia, cognitive impairment or dementia. The Cambridge Psychology Research Ethics Committee approved the study, and all participants gave written informed consent before participating and were compensated. Table 1 summarises demographics, excluding outliers, broken down by time of day (morning versus evening), experiment format (video versus non-video), and age (younger adult versus older adult).

**Table 1.** Demographics of Younger and Older Adults in Video vs. non-Video Format (means & standard deviations).

| Condition | Age group | N | Age (SD) | Gender | Time of day | Years of education (SD) |
| --- | --- | --- | --- | --- | --- | --- |
| **Video (Supervised)** | YA | 73 | 24.8 (3.74) | 36 F, 37 M | 35 Morning<br>38 Evening | 16.5 (2.46) |
| **Video (Supervised)** | OA | 71 | 65.4 (4.41) | 36 F, 35 M | 36 Morning<br>35 Evening | 15.8 (3.46) |
| **Non-Video (Unsupervised)** | YA | 46 | 26.2 (3.40) | 22 F, 23 M, 1 O | 25 Morning<br>21 Evening | 16.1 (3.26) |
| **Non-Video (Unsupervised)** | OA | 42 | 66.0 (5.28) | 21 F, 21 M | 21 Morning<br>21 Evening | 15.0 (3.22) |

Total N = 232. M = Males; F = Females; O = Other. YA= Younger Adults; OA = Older Adults.
Standard deviations are reported in parentheses.

### Design and procedure

The experiment was constructed using PsychoPy [21] and carried out online (pavlovia.org). In the video condition, participants completed the memory task while on a video call with the researcher, ensuring attention was maintained throughout the task. In the non-video condition, no researcher was present during the task, however, attention checks were implemented throughout the learning and recall phases in the non-video condition to promote sustained attention. Testing sessions were divided into morning sessions (7:00 a.m. to 10:00 a.m.) and evening sessions (7:00 p.m. to 10:00 p.m.). Participants were able to choose from a morning or evening session. The memory task was adapted from a prior study which used a continuous memory paradigm to investigate the impact of schematic memory on episodic retrieval over time [22].

#### Materials

Stimuli for the memory task consisted of 120 everyday objects obtained from stimuli image sets [23–25] and Google image search. Each object was superimposed on a light grey background (Fig 1). Locations for each object were randomly selected from a circular (0-360 degree) parameter space. Stimulus generation was conducted once, and all participants learned the same displays. Order presentation of displays was randomised across blocks.

**Fig 1.**
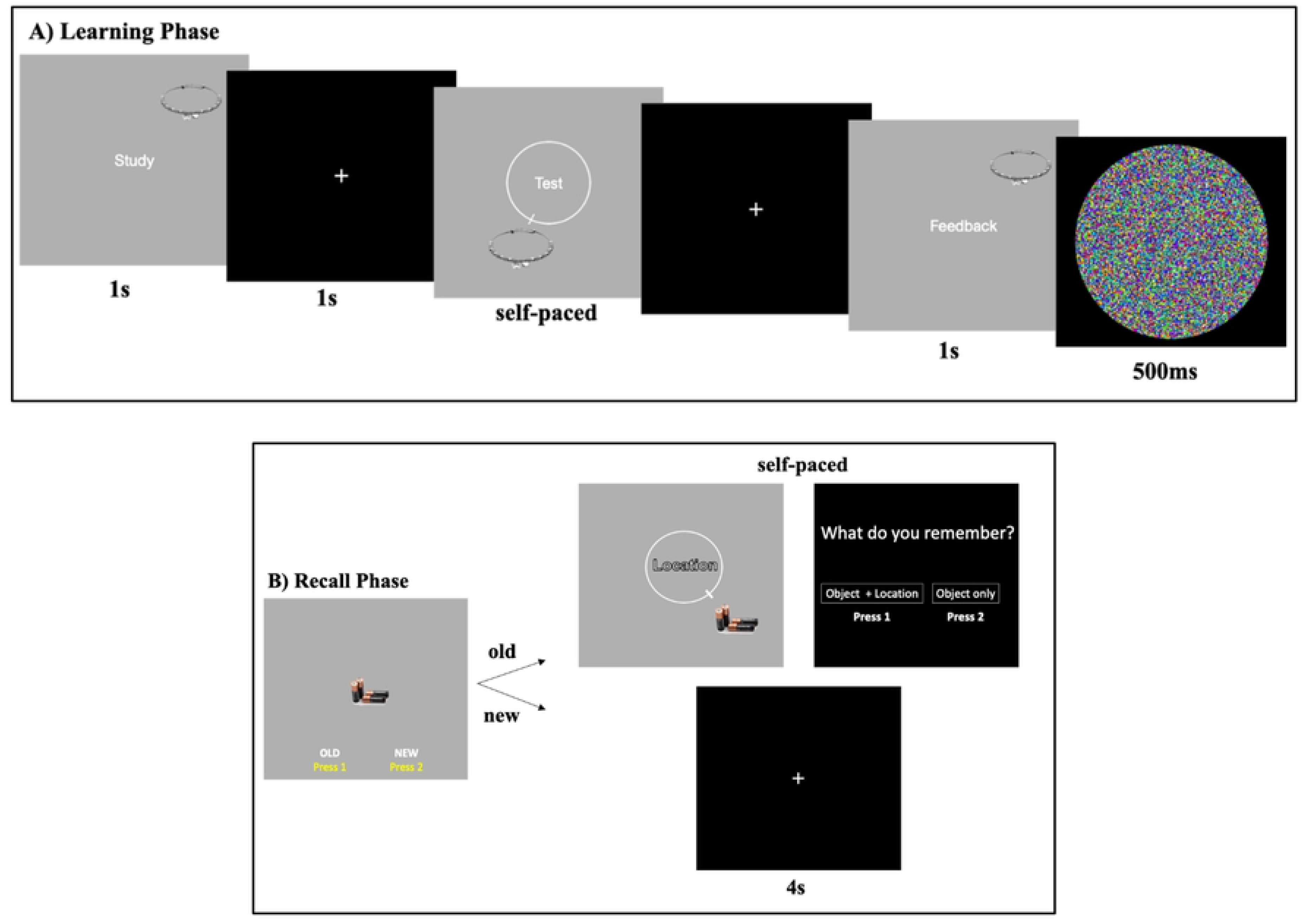
Example Trials from (A) Learning and (B) Recall Phase.

#### Learning phase

Prior to the start of the task, participants were instructed that they would learn and be tested on the locations of objects displayed at different positions around a circular dial. The researcher read aloud the instructions for the task, and participants then completed a short practice learning and recall phase to become familiarised with the task and ask any clarifying questions before the start of the first block.

In each learning phase, participants learned a series of displays consisting of one real-world object pseudo-randomly placed around a circle overlaid on a grey background. Each display was presented for one second, accompanied by the word “Study” in the centre of the display. Participants were instructed to remember both the identity and the location (target to-be-remembered location) in each display. Following a brief inter-trial interval (one-second fixation cross), the same object was displayed in a random location around a circular dial. The word “Test” was presented in the centre of the dial to indicate that the participants’ memory for the item’s location was now going to be tested. Participants were instructed to move the studied object around the dial using the left and right arrow keys to as close as possible to the location they previously saw within the display and press the spacebar to submit their answers. Responses were self-paced. However, if no response was made after 6 seconds, the colour of the word “Test” changed from white to red to encourage the participant to respond in a timely manner.

Following the relocation, participants were presented for one second with the same object in its correct target location, accompanied by the word “Feedback” in the centre of the display to remind the participant of the object’s target location. Each trial component was separated by a one-second fixation period. To indicate the end of learning the location of an object, a multicoloured circle mark image briefly appeared (500 ms) on the screen. The 120 stimuli were presented over three blocks. Each block consisted of a learning phase (40 objects), followed by an immediate retrieval test of half of the current block stimuli (20 objects) plus eight new items. Assignment of objects to learning phase blocks and order of presentation were randomised across participants.

#### Recall phase

Each learning phase was followed by an immediate recall phase. Participants were presented either with an object studied in the prior learning phase or a new object not shown in the learning phase. In each trial of the recall phase, the participants’ task was first to identify whether the object was old, meaning previously studied (pressing one), or if the object was new, meaning not shown in the previous learning phase (pressing two). For objects chosen as old, regardless of whether the object was or was not previously studied, the object was displayed in a random location around a circular dial. The word location was presented in the centre of the dial to indicate that participants’ memory for the item’s location would then be tested. Participants were instructed to move the object as close as possible to the location where they previously saw it within the display during the first task (learning phase). Once a response was made, participants were then asked to indicate if they remembered both the object and its associated location (Object + Location) from the learning phase by pressing 1, or if they just remembered the object but not its associated location (Object only), to press 2. For objects chosen as New, location response and subjective judgments were not made. However, to account for similar timing between old and new responses and deter participants from pressing new to advance the study, a fixation cross appeared in the centre of the screen, and there was a short delay period (4 seconds) before the next trial began. Participants were tested on half (20) old objects from the previous learning phase (targets), and eight new items that were not presented in the prior learning phase (foils). Objects to be tested in the immediate recall were chosen at random, and the presentation of old/new objects was randomised.

At the end of the memory task, participants were redirected to a short questionnaire that contained the Pittsburgh Sleep Quality Index (PSQI), a 19-item self-rated questionnaire assessing sleep quantity (e.g., number of hours of sleep), quality (subjectively measured), and disturbances over the course of a month [26]. In addition, participants were also given the Morningness-Eveningness Questionnaire (MEQ), a 19-item self-reported assessment of questions related to preferred wake/sleep times, timing of physical and mental activities and alertness throughout the day [10]. Answers to these questions were used to examine the effects of circadian preferences on cognitive performance in the memory task and served as a comparison to the time the task was performed.

## Modelling and data analysis

### Recognition memory

Memory performance was first assessed by calculating the proportion of “old” responses to previously studied objects (targets) and the proportion of “old” responses to new objects (foils) not presented during the learning phase. A standard recognition score was calculated by subtracting the proportion of “old” responses to foils (false alarm rate) from the proportion of “old” responses to targets (hit rate). This score reflects participants’ ability to correctly identify old items presented during the learning phase while rejecting new objects [27,28]. A three-way ANOVA examined the influence of recognition score as a function of age group (older vs. younger), time of day (morning vs. evening), and experiment format (video vs. non-video) on recognition memory.

### Non-parametric subject-level outcomes

Retrieval error was calculated for each trial as the angular deviation between the object’s target location during the learning phase and the participants’ response during the immediate recall phase. The difference between the target and the response provides the absolute error (0-180 degrees). Average recall error was calculated for each subject, and a three-way ANOVA examined the influence of age group, time of day, and experiment format on mean retrieval error.

### Morningness-Eveningness Questionnaire (MEQ)

Each question from the MEQ was scored and summed, providing a score ranging from 16 to 86. Scores of 41 and below indicate “evening types”, scores of 59 and above indicate “morning types”, and scores between 42 and 58 indicate “intermediate types.” An ANOVA examined whether MEQ scores differed as a function of age group and time of day. In addition, Bayes factor analyses were performed in JASP [29] using model averaging across matched models to quantify the strength of evidence for individual main effects and interaction terms. For non-significant effects, Exclusion Bayes factors (BF_excl_) >3 was considered moderate evidence in favour of the null hypothesis [30,31]. For significant effects, Inclusion Bayes factors (BF_incl_) were reported to quantify evidence for the alternative hypothesis. In addition, to further assess correspondence between chronotype and time of testing (morning vs evening), a median split was applied to MEQ scores (high vs. low MEQ scores), and participants were classified as either “matched” (MEQ score aligned with time of testing) or “mismatched” (MEQ score did not align with testing time).

### Bayesian hierarchical mixture modelling

In contrast to prior memory precision studies, which primarily used a two-step, non-hierarchical modelling method for parameter estimation, the need in the present study to simultaneously examine multiple categorical (e.g., age, time of day, and experiment format) and continuous (e.g., MEQ) predictors requires an advanced, hierarchical approach. Traditionally, parameters are first estimated for each subject in a particular condition (e.g., age group) and then further analysed with subsequent tests (t-tests, ANOVA, linear regression) to make inferences for how those parameters vary as a function of other conditions (e.g., time of day, experiment format). This approach is limited, however, in that it fails to account for parameter uncertainty in hierarchically related data (e.g., related age or demographic groups, experimental format groups, and/or groups of conditions such as location or time of day) [32].

In the current experiment, response errors were estimated using a two-parameter Bayesian hierarchical mixture model to investigate two possible sources of retrieval errors: variability in memory precision and memory retrieval failures (e.g., guessing). The model comprised a mixture of two von Mises distributions, which were specified to ensure that both covered the same range of responses. The first component consisted of a von Mises distribution (the circular equivalent of a Gaussian distribution) centred on the target location, which reflects variability in the precision of remembered locations. Parameter estimates obtained from the target von Mises distribution provided a measure of the precision of target retrieval, estimated from the parameter κ₁ (precision). The second component was a von Mises distribution with precision zero (equivalent to a uniform distribution over a circular space), representing failures in memory retrieval, in which responses are random guesses. The mixture weight associated with this component is θ₁ (probability of memory-based responding), which is transformed into probabilities (Pₑ)(θ₁) and (P_guess_)(1-θ₁). Each distribution yields two classes of parameters: distributional parameters (mean, μ, and precision, κ) and mixing proportions (*theta,* θ).

Bayesian Hierarchical Mixture Modelling was implemented using the brms package [33,34] with a Stan backend in R (version 4.3.1; R Core Team, 2023).This hierarchical model yields parameters, namely, precision (κ₁) and the probability of memory-based responding (θ₁), analogous to those obtained from standard (non-hierarchical) mixture models [35,36]. However, unlike traditional approaches that fit separate models with and without a guessing component, these parameters are estimated simultaneously within a single hierarchical framework. In addition, Bayesian Hierarchical Mixture Modelling allows for parameters to be estimated simultaneously across all conditions (e.g., age, time of day, experiment format) rather than separately for each group or condition, requiring subsequent statistical comparisons (t-test or ANOVAs), which can constrain parameter estimations [32]. In addition, non-hierarchical models can be less stable with fewer trials. This is particularly relevant in this experiment, as modelling is based on correctly identified objects; therefore, the number of trials varies across participants. In all, hierarchical modelling estimates parameters across all subjects and conditions simultaneously and allows for estimation of individual differences, producing more robust and reliable parameter estimates.

#### Primary model: time of day, age, and experiment format

Precision (κ₁) and memory probability (θ₁) were modelled as functions of age group (younger vs. older), time of day (morning vs. evening), and experiment format (video vs. non-video). The model included main effects of all predictors, as well as interaction terms between age group and each experimental factor (age group x time of day; age group x format). Both κ₁ and θ₁ included varying intercepts by participant (SubjectID), allowing for assessment of individual differences in memory precision and response probability.

#### Secondary model: chronotype (MEQ), age, and time of day

To assess whether the time at which a participant completed the memory task corresponded to their optimal or preferred time (synchrony effect) and whether this influenced their performance, a second Bayesian hierarchical mixture model was fit to participants’ error data. The model included chronotype (MEQ, z-scored), time of day (morning vs evening), and age group (older adult vs younger adult), along with all two- and three-way interactions.

#### Specification of priors

Selection of priors is essential to ensure the model produces reliable parameter estimates.

Priors were chosen based on recommendations from prior visual working memory tasks that used Bayesian hierarchical mixture modelling [37]. The same priors were used for both models, except that only the predictor variables differed (experiment format in the primary model and MEQ in the secondary model).

Before fitting the hierarchical model, prior predictive checks were implemented to ensure that the chosen priors would accurately inform the model and fit the observed data well. In particular, prior sensitivity analyses were conducted in which data was simulated from the model using either uninformative (A, weak), informative (B, mild), or conservative (C, tight) priors.

Prior-predictive distributions were visually compared, and informative priors (Fig 2B) were chosen for the model (described below) as they best predicted expected data allowing for a plausible range of precision and memory probability values (Fig 2).

**Fig 2.**
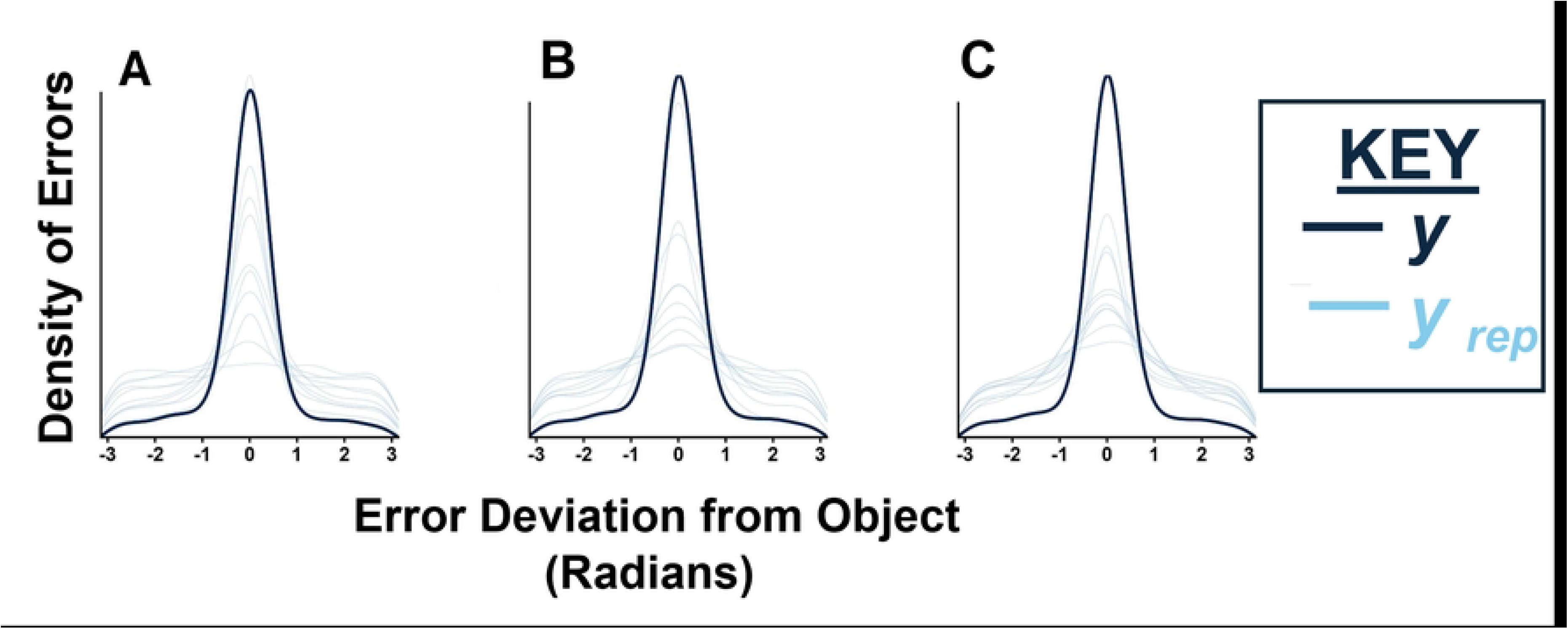
Posterior Predictive plot for uninformative (A), informative (B), or conservative (C) priors.

Specifically, only the precision of the target distribution (κ₁) and the probability of memory-based responding (θ₁) were freely estimated, whereas remaining parameters were either fixed or tightly constrained via priors. The means (μ₁, μ₂) of both von Mises distributions were fixed at zero using Constant (0) priors following the assumption that responses are centered on the target location. The precision parameter is modelled on the log scale. Therefore, for the precision parameter (κ₁), the intercept was assigned a weakly informative Normal (0, 1) prior, encompassing a broad but plausible range of baseline precision values. The regression coefficient for (κ₁) was assigned Normal (0, 0.5) priors, allowing precision to either increase or decrease across conditions. The mixing proportion (θ₁), which represents the probability of memory-based responding, is modelled on the logit scale. Therefore, the intercept was assigned a Normal (0, 1) prior, allowing for a broad but reasonable range of baseline memory probabilities. The regression coefficients for (θ₁) were assigned a more regularising Normal (0, 0.3) encompassing a plausible range of changes in probabilities.

For the second distribution, the precision parameter (κ₂) was fixed to a very small value (log κ₂ = -10), to approximate a uniform distribution over the circular space, consistent with random guessing. The mixing proportion (θ₂) was not estimated directly (fixed to 0 by default) and corresponds to 1-θ₁.

The model included random intercepts for both κ₁ and θ₁ to estimate the influence of individual differences in precision and memory probability. Parameters were estimated using four Markov chain Monte Carlo (MCMC) chains, each with 500 warm-up samples, 2,200 iterations, and 6800 post-warm-up draws. Each model was fit to 11,118 observations.

#### Assessment of model fit and parameters

Model fit for both models was first assessed by visually inspecting posterior predictive plots of error distributions, examining the extent to which the observed data aligned with the model’s posterior-predictive response distributions. In addition, convergence of the MCMC chains was assessed by checking for divergent transitions during sampling and by verifying that R^ values did not exceed 1.05.

Regression coefficients summarised the effects of predictors (e.g., age, time of day, and experiment format (primary model) or MEQ score (secondary model)) on model parameters.

Multilevel hyperparameters were examined to assess the influence of individual differences on memory precision (κ₁) and the probability of memory-based responding (θ₁). Uncertainty in parameter estimates was quantified using 95% Bayesian credible intervals, with intervals including zero indicating limited evidence of an effect.

Posterior estimates of model parameters were obtained on the transformed (linear-predictor) scale (log for κ₁ and logit for θ₁) and converted using the inverse link functions (e.g., exponentiation for κ₁ and logistic transformation for θ₁). For interpretability, κ was further transformed into circular standard deviation of the von Mises distribution (degrees), reflecting memory imprecision. θ₁ was transformed and expressed as the probability of memory-based responding (Pₑ).

For the first model, which contained only categorical predictors, condition-specific posterior estimates were obtained by reconstructing linear predictors from the model coefficients. For the second model that included a continuous predictor (MEQ), posterior predictions were generated over a continuous range of MEQ values (z-scores from −2.5 to 2.5). For both models, summary estimates were computed as posterior medians with 95% credible intervals.

In addition, to verify if the current sample size (*N* = 232) was large enough to estimate the effects, the hierarchical mixture model was fit to subsets of the data (*N =* 50, 100, 150, 200, 232), and posterior distributions for the parameters of interest (e.g., format, time of day, and age group interactions) were examined to assess how they change as sample size (*N*) increases (See Supplementary).

#### Posterior evidence evaluation using ROPE-HDI decision

The Region of Practical Equivalence (ROPE) and HDI (Highest Density Interval) decision rule were applied to evaluate the strength of evidence for each interaction parameter in the model. In this approach, ROPEs are defined as a “range of parameter values that are equivalent to the null value for practical proposes” [38,39]. The HDI of a posterior distribution is defined as the interval that contains the 95% most credible values of a parameter, such that points that fall within this interval are considered more credible (higher density) than those that fall out of this interval (lower density). Together, the HDI + ROPE decision rule can be used to assess whether effects are meaningfully different from zero such that if the entire 95% HDI lies outside of the ROPE (95% of the most credible values of parameters are not equivalent to the null value), this demonstrates evidence of a meaningful effect (reject the null). In contrast, if the 95% HDI falls completely inside the ROPE, the credible values are practically equivalent to the null, providing evidence the effect is not meaningful (accept the null). Finally, if the HDI partially overlaps with the ROPE, the conclusion remains undecided as some credible values are practically equivalent to the null, while others are not.

ROPE’s were defined for each parameter (precision κ₁, memory probability θ₁) based on prior research, which typically reports medium-sized effects for age-related differences in memory precision and retrieval success [7, 40], as well as for time-of-day effects on cognitive performance [11,14,41]. For example, age-related reductions in memory precision correspond to differences of approximately 0.15-0.40 (log scale), therefore we defined the ROPE for precision parameters to be ±0.10 log units, representing a conservative threshold for what we would consider to be a practically negligible effect (smallest effect size of interest). With respect to memory probability, age related and time of day effects on memory performance have been observed to fall within the range of 0.25-0.60 (logit scale), so the memory probability threshold was set to ±0.35 such that values outside this range would not represent a meaningful difference. Interaction parameters were evaluated using the HDI-ROPE decision rule, examining whether the 95% HDI lay entirely inside, outside, or are overlapping with the defined ROPE. In addition, the strength of evidence was quantified providing a continuous measure, by calculating the posterior probability that each interaction parameter lay inside or outside the ROPE. All data and code used to run experiments and conduct analyses, including hierarchical modelling code, are available on the Git Hub repository.

## Results

### Recognition memory

A three-way ANOVA revealed a significant main effect of experiment format (F(1, 224) = 4.68, *p* = 0.032) on recognition scores with video-based sessions (*M* = 0.77) yielding higher performance than non-video sessions (*M* = 0.73) (see Fig 3). However, Bayes factor indicated only weak evidence for a difference in format (*BF_incl_* = 1.32). There was no evidence for a main effect of age group (F(1, 224) = 1.99, *p* = 0.16, *BF_excl_* = 1.40), with older adults (*M* = 0.78) performing similarly to younger adults (*M* = 0.74). There was no main effect of time of day (F(1, 224) = 0.05, *p* = 0.83, *BF_excl_*= 6.74). There was also no interaction between age group and time of day (F(1, 224) = 0.12, *p* = 0.729, *BF_excl_* = 5.14).

**Fig 3.**
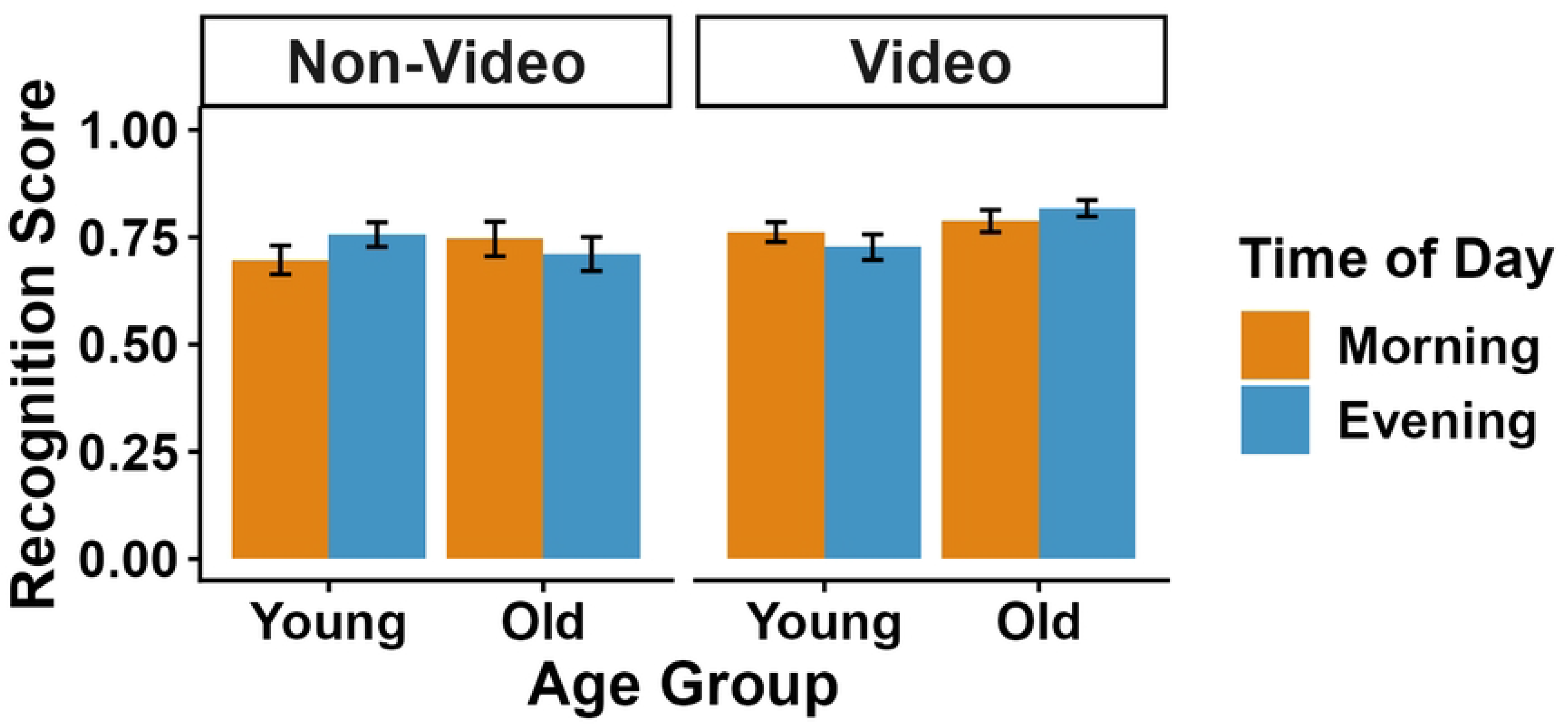
Recognition Performance by Age Group, Time of Day, and Experimental Format. Bars represent mean recognition scores. Error bars indicate ±1 standard error of the mean.

Although the three-way interaction between age group, time of day, and experiment format approached significance (F(1, 224) = 3.52, *p* = 0.062), Bayes analysis proved inconclusive (*BF_excl_* = 0.73), indicating a lack of sufficient evidence to confirm the presence or absence of an interaction. Fig 3 depicts recognition scores broken down by experiment format (video vs. non-video), age group (older vs younger), and time of day (morning vs. evening).

### Learning and recall error

To assess learning and immediate recall accuracy, Mean Absolute Error (MAE) was calculated across all trials for each participant during both the learning and immediate recall phases. MAE values were computed separately for younger and older adults in each time-of-day condition and for both video and non-video testing sessions.

A linear regression analysis revealed that Learning MAE significantly predicted Recall MAE (*β* = 1.79, *p* < 0.001), indicating that participants who performed better during learning also tended to perform better during immediate recall. Experimental format was also a significant predictor, with the video condition associated with lower Recall MAE (*β* = -6.09, *p* = 0.016), suggesting improved performance or reduced recall error in the video condition. There were no significant effects of age group (*β* = 3.66, *p* = 0.138) or time of day (*β* = 1.28, *p* = 0.598) on recall MAE. The overall model accounted for a modest proportion of variance in Recall MAE (*R*² = 0.15).

### Bayesian hierarchical modelling results of model one (time of day, age, format)

#### Assessment of model fit

Fig 4 illustrates a posterior predictive plot [37] of the model one fit, comparing 10 model-predicted error distributions (*y rep*; light blue lines) with the observed data distribution (*y*; dark blue line). The posterior predictive distribution of errors indicated a good fit of the model to the data, as evidenced by the overlap between the observed and simulated data. The Markov Chain Monte Carlo (MCMC) chains showed good convergence across all parameters (R^ = 1.00), and effective sample sizes (Bulk ESS and Tail ESS) were high (>2000 for most parameters), indicating stable posterior estimation.

**Fig 4.**
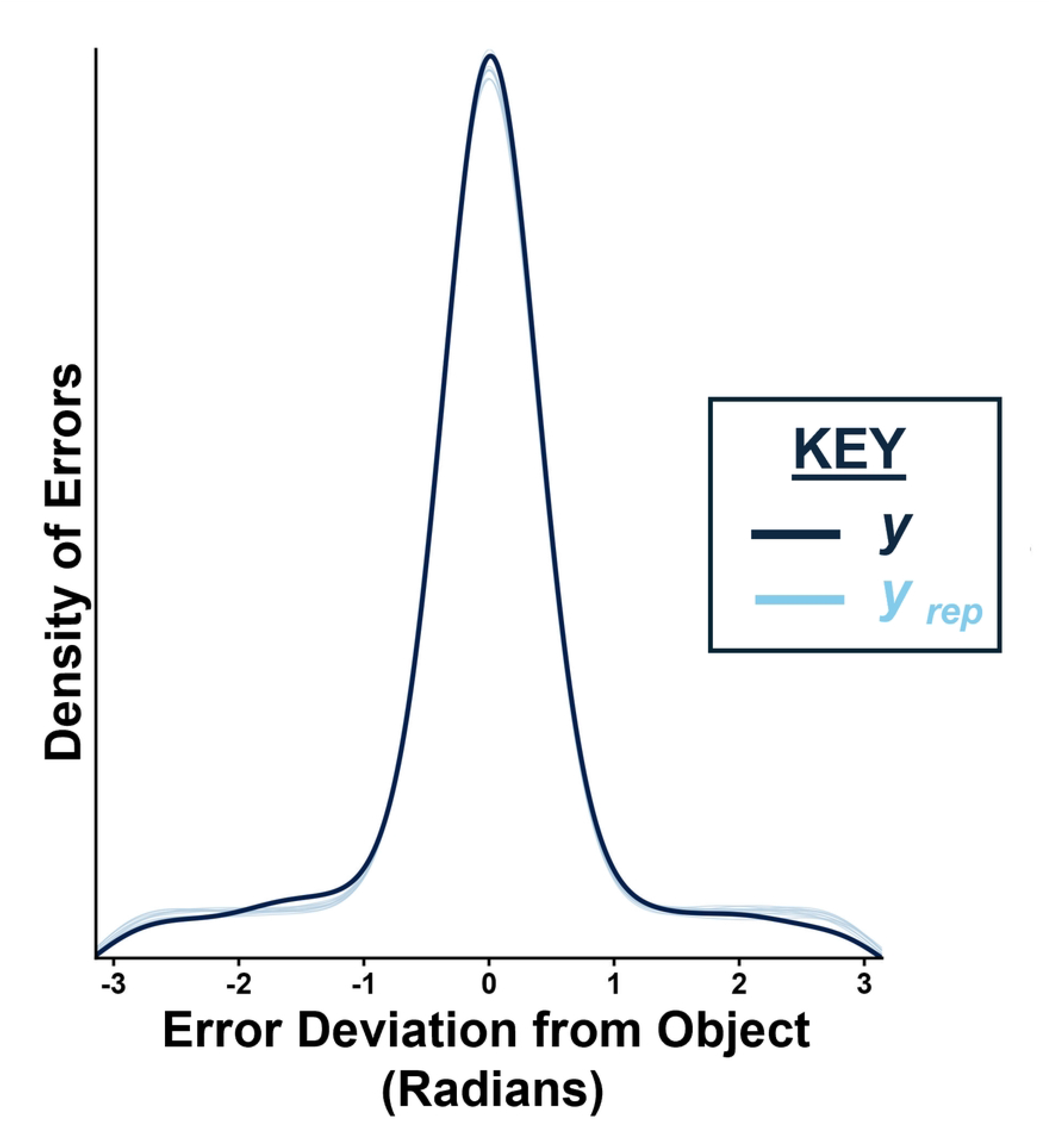
Posterior Predictive Plot of Model One Fit (Predicted (y rep) vs. Actual (y) Data Distribution)

#### Model Results

##### Memory precision (κ₁)

Baseline memory precision for the reference condition (younger adults in the video format during the day) was estimated as κ₁ = 2.99, 95% CI [2.79, 3.19]. There was evidence of a negative effect of age group on precision (β = -0.27, 95% CI [-0.53, 0.01]), indicating that older adults tended to exhibit lower memory precision than younger adults, although the effect was moderate and included zero (See Fig 5A). There was no evidence for an effect of testing session format (β = −0.01, 95% CI [-0.26, 0.25]) or time of day (β = −0.08, 95% CI [-0.33, 0.16]) on precision. Additionally, there was no evidence of an interaction between age group and format (β = 0.08, 95% CI [-0.28, 0.43]) or between age group and time of day (β = 0.07, 95% CI [-0.27, 0.40]).

**Fig 5.**
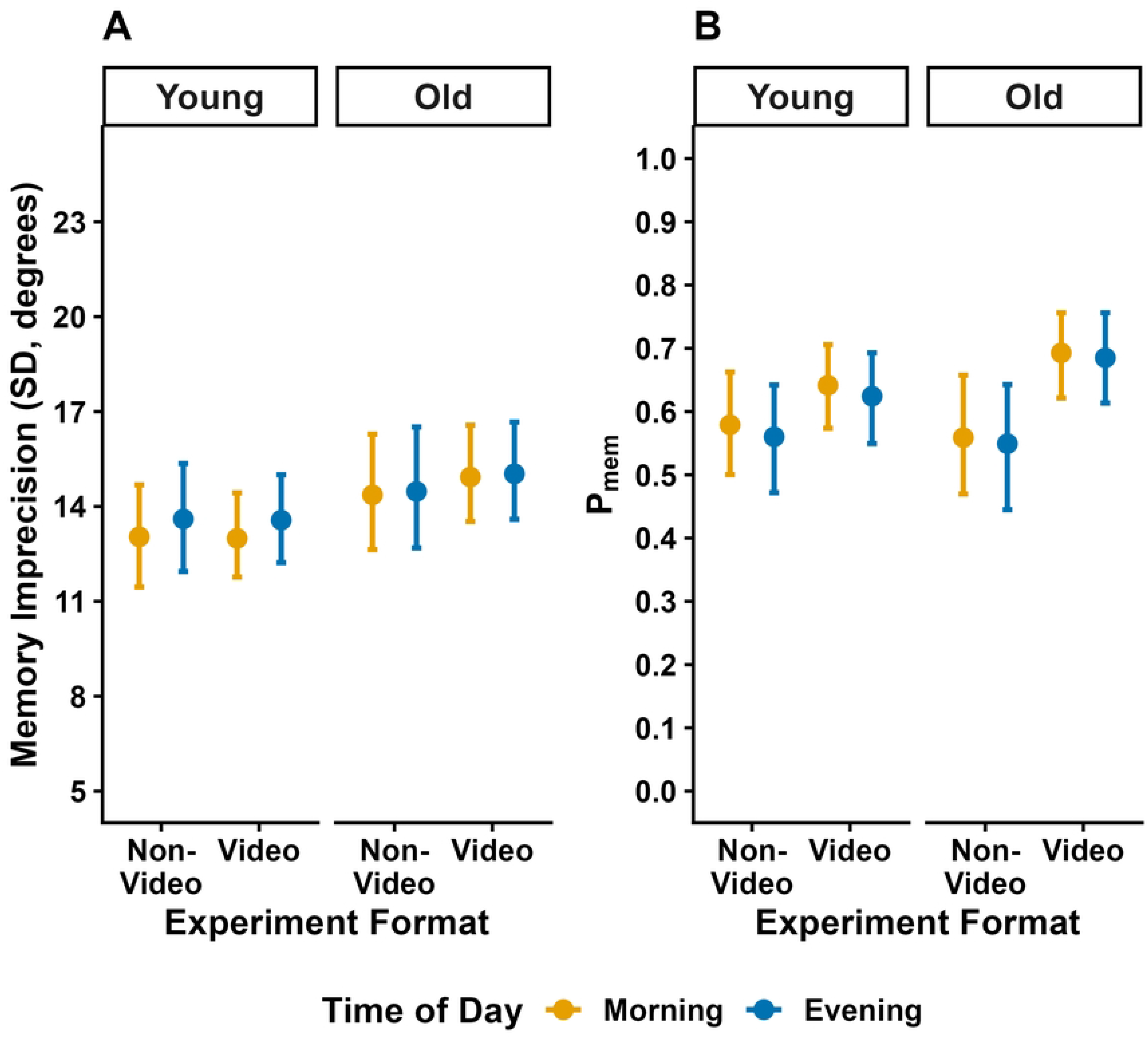
Parameter Estimates of Imprecision of Memory (A) and Memory-based Responding (B). The figure shows posterior medians (points) and 95% credible intervals (lines) for (A) memory imprecision (standard deviation in degrees) and (B) the probability of memory-based responding (θ₁; Pₑ) as a function of age group (older vs younger), format (video vs. non-video), and time of day (morning vs. evening).

##### Probability of memory-based response (θ₁)

Baseline probability of memory-based responding for the reference condition was estimated as θ1 = 0.58, 95% CI [0.29, 0.87]. There was no evidence for an effect of age group (β=0.23, 95% CI [-0.11, 0.58]), testing format (β=−0.26, 95% CI [-0.59, 0.06]), or time of day (β=−0.07, 95% CI [-0.40, 0.25]) on the probability of recalling an object from memory (See Fig 5B). In addition, there was no evidence for an interaction between age group and format (β = −0.31, 95% CI [-0.71, 0.10]) or between age group and time-of-day (β = 0.04, 95% CI [-0.35, 0.43]).

##### Between-subject variability

The model indicated substantial between-subject variability in both memory precision (SD = 0.53, 95% CI [0.44, 0.62]) and probability of memory-based responding (SD = 1.20, 95% CI [1.06, 1.35]), suggesting meaningful individual differences across participants.

##### Visualisation of posterior estimates

Visualisation of posterior estimates showed substantial overlap in credible intervals and posterior distributions across conditions (time of day and experiment format), consistent with the absence of strong effects. In contrast, age differences were evident for memory precision, with older adults exhibiting reduced precision relative to younger adults (See Fig 5).

### Age, time of day, MEQ

Independent of memory performance, we examined how MEQ score (reflecting chronotype) aligned with time of testing and how this differed with age. A two-way ANOVA on MEQ scores revealed a significant main effect of age group (*F*(1, 228) = 59.86, *p* < 0.001, *BF_incl_* = 1.99 x 10^10^), such that older adults’ mean MEQ scores were higher, indicating a morning chronotype (*M* = 57.9, *SE* = 0.81), compared to younger adults’ MEQ scores (*M* = 49.11, *SE* = 0.79). There was a main effect of time of day, driven by those in the morning condition, who had higher MEQ scores (M = 55.20, *SE* = 0.80) than those in the evening condition (*M* = 51.77, *SE* = 0.82); (*F*(1, 228) = 9.19, *p* = 0.003, *BF_incl_* = 9.85). However, there was no significant interaction between time of day and age on MEQ scores (*F*(1, 228) = 0.018, *p* = 0.894, *BF_excl_* = 4.68), indicating moderate evidence in favour of the null hypothesis (see Fig 6). As shown by median split analysis, some participants were tested at their preferred time of day (matched; *N* = 134, Older = 64, Younger = 70) and others at their non-preferred time (mismatched; *N* = 98, Older = 49, Younger = 49).

**Fig 6.**
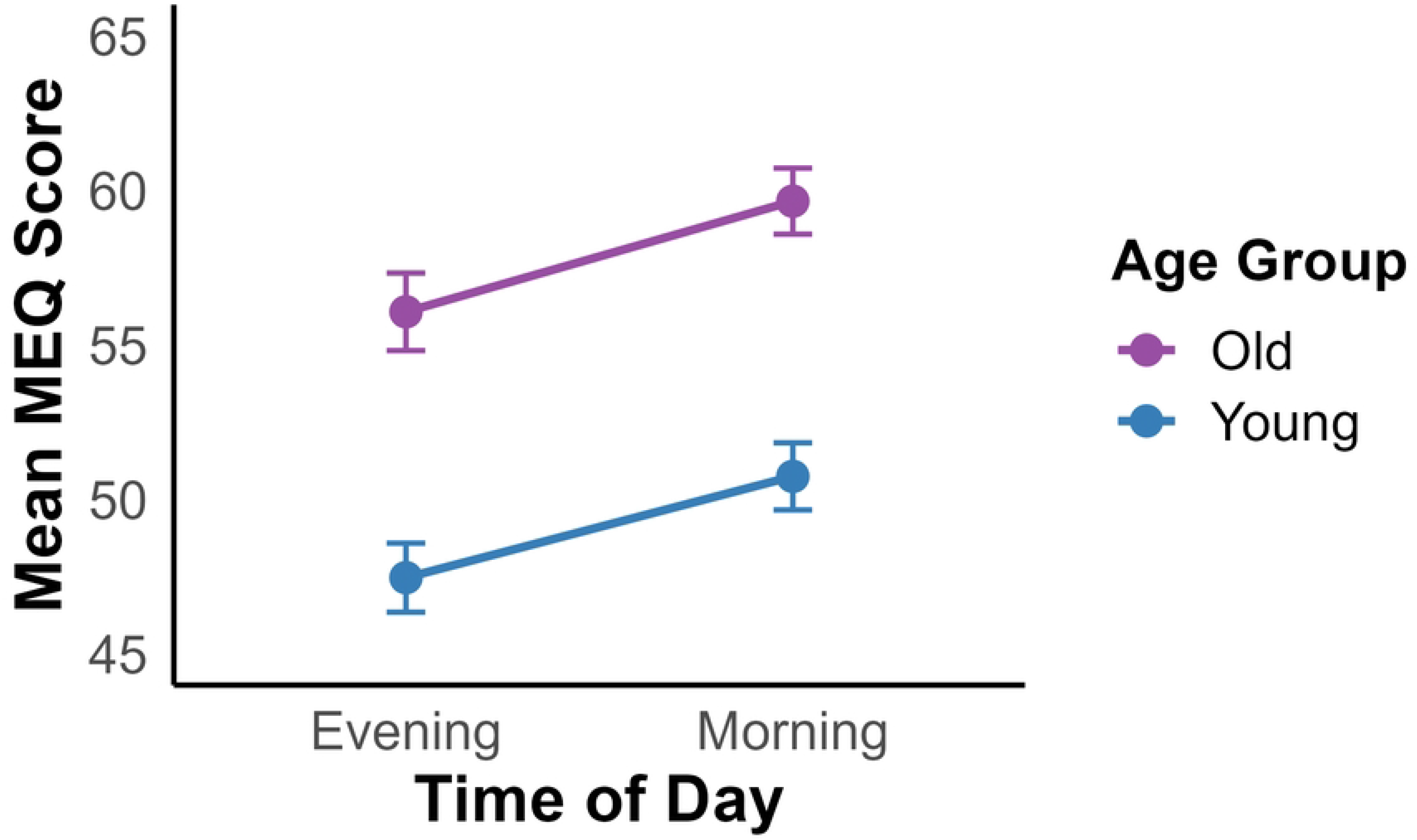
Mean Morningness–Eveningness Questionnaire (MEQ) scores by Age Group and Time of Day. MEQ scores range from 16 to 86, with higher values indicating a stronger morning chronotype. Error bars represent ±1 standard error.

### Bayesian hierarchical modelling results of model two (MEQ, age, and time of day)

#### Assessment of model fit

Similar to model one (age, time of day, format), the posterior predictive distribution of errors in model two (MEQ, age, time of day) indicated a good fit to the data, as evidenced by the overlap between the observed and simulated data (see Fig 7). The Markov Chain Monte Carlo (MCMC) chains showed good convergence across all parameters (R^ = 1.00), and effective sample sizes (Bulk ESS and Tail ESS) were high (>2000 for most parameters), indicating stable posterior estimation.

**Fig 7.**
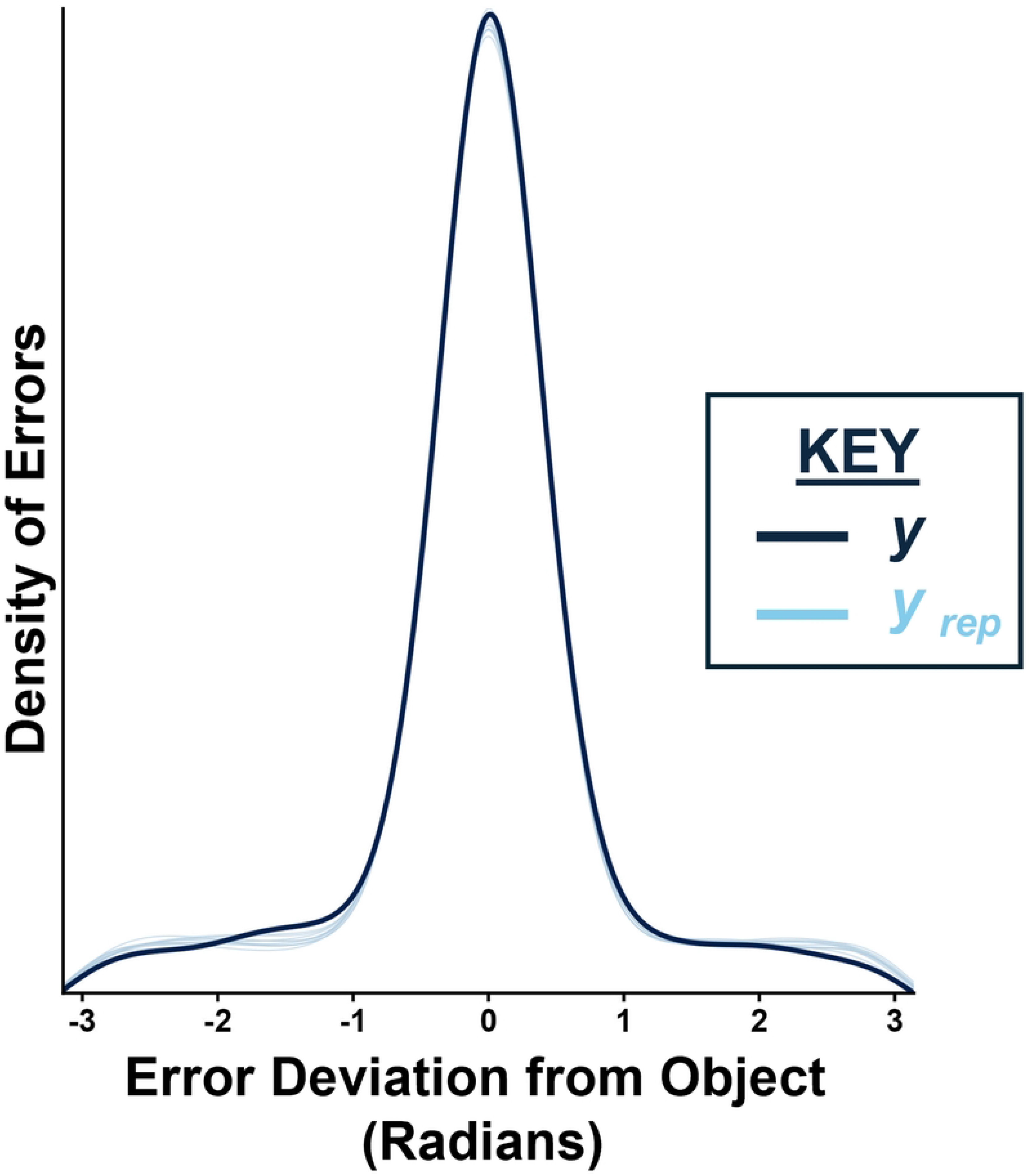
Posterior Predictive Plot of Model Fit of Predicted vs. Actual Data Distribution for Model Two.

#### Model results

##### Memory precision (κ₁)

The Bayesian hierarchical model indicated little evidence that memory precision (κ₁) varied with chronotype, time of day, or their interaction (See Fig 8A). Importantly, the interaction between MEQ and time of day, capturing the predicted synchrony effect, was small and uncertain (β = −0.13, 95% CI [−0.41, 0.14]). In addition, there was no evidence that this relationship differed by age group, as the three-way interaction between age group, time of day, and MEQ was small (β = 0.16, 95% CI [−0.21, 0.51]), indicating no reliable age-related modulation of the synchrony effect.

**Fig 8.**
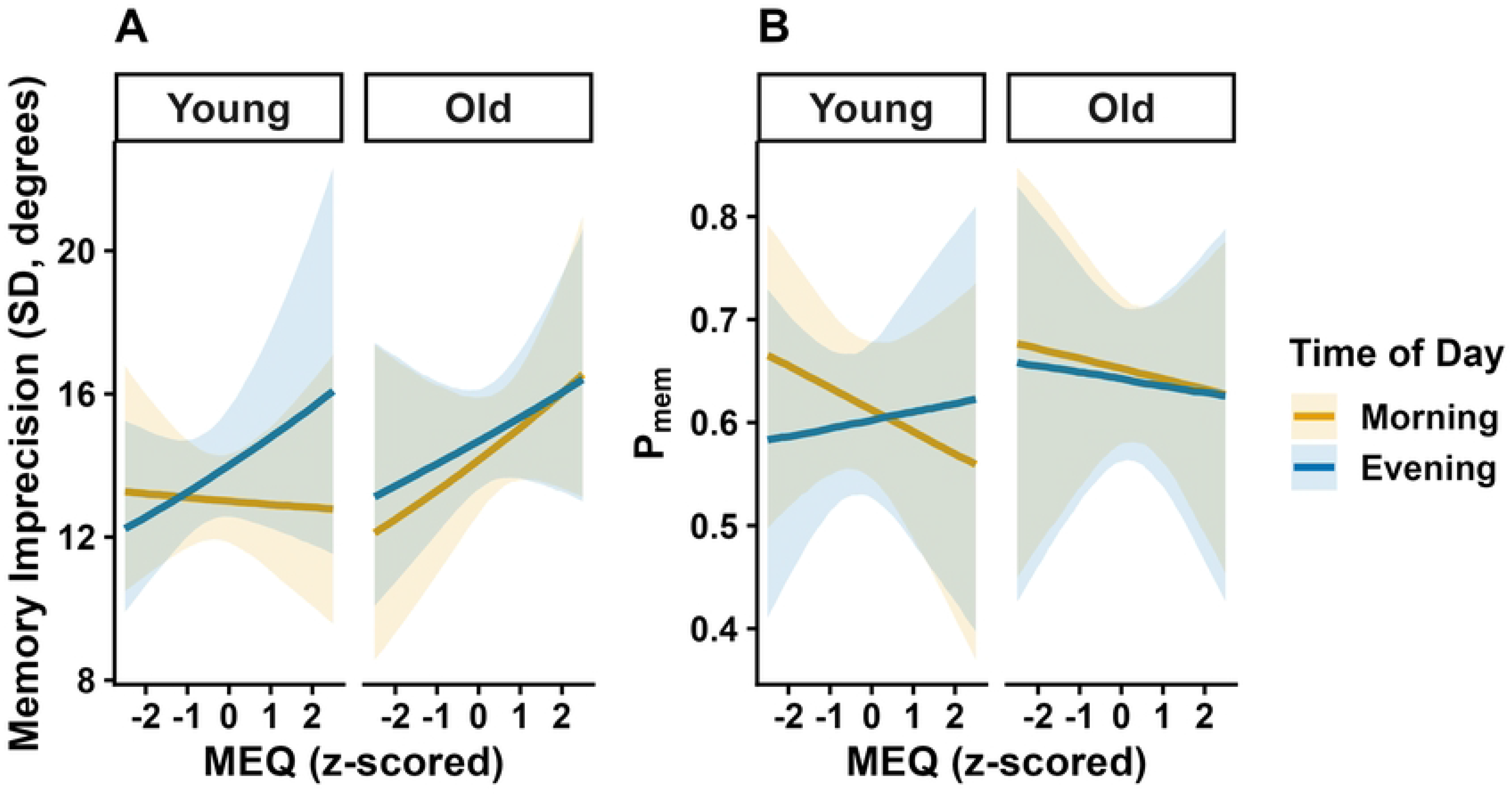
Model-derived Effects of Chronotype, Age and Time of Day on Memory Performance. Estimated memory imprecision (circular SD in degrees, derived from κ; (A), and probability of memory-based responding (Pₑ) (B). Lines represent posterior medians, with shaded regions indicating 95% credible intervals. Predictions are plotted across the range of MEQ (z-scored), separately by age group and time-of-day condition.

##### Memory Probability (θ₁)

A similar pattern was observed for the probability of memory-based responding (θ₁) (See Fig 8B). The interaction between MEQ and time of day was small and uncertain (β = 0.13, 95% CI [−0.20, 0.46]), providing no evidence that participants were more likely to retrieve memories when tested at their optimal time of day. The three-way interaction with the age group was also not reliably different from zero (β = −0.11, 95% CI [−0.52, 0.31]), such that the synchrony effect did not vary between younger and older adults.

Posterior predictions show substantial overlap in credible intervals across time of day for both younger and older adults (Fig 8B).

#### Posterior evidence evaluation using ROPE-HDI decision

##### Memory probability (θ₁)

As depicted in Fig 9, the posterior distribution for memory probability for age group by time of day interaction (Fig 9A), was centered near zero (M = 0.04, 95% HDI ([-0.36, 0.43]). Visually, overlap between the HDI and ROPE suggested uncertainty, however, when examining the posterior probability, a majority fell within the range of practical equivalence (*p*_ROPE_ 91.4%) and only 8.6% exceeded the ROPE threshold for a meaningful effect.

**Fig 9.**
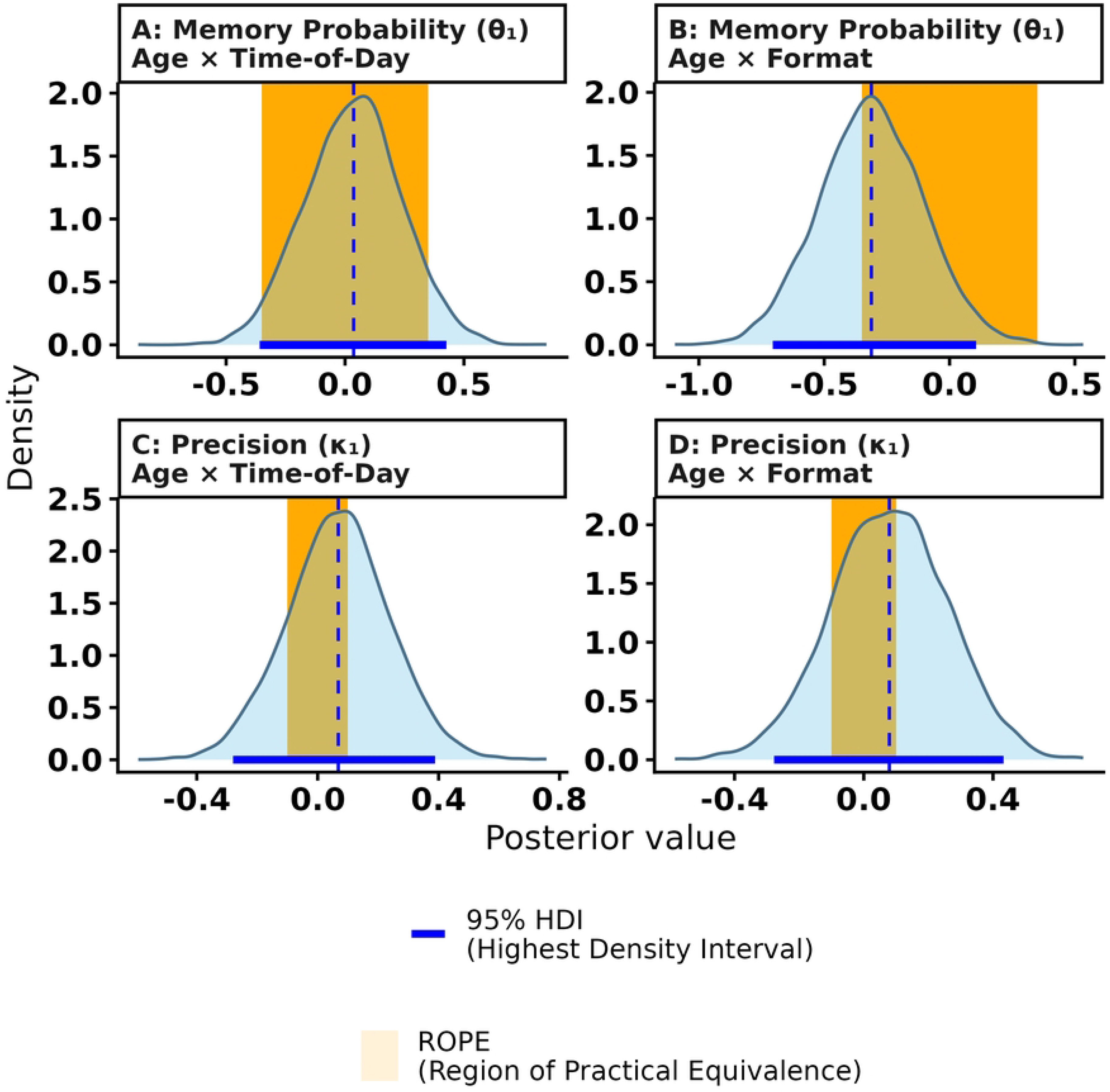
Posterior Density Plots for Age Group by Time of Day and Age Group by Format. Interactions are depicted on memory probability (θ₁) and memory precision (κ₁). Each panel depicts the posterior distribution (light blue), the 95% Highest Density Interval (HDI; solid blue line), the posterior mean (dashed line). The Region of Practical Equivalence (ROPE), defined as ±0.35 for θ₁ and ±0.10 for κ₁, is displayed by the orange shaded region.

The age group by experiment format interaction (Fig 9B) exhibited a negative posterior mean (M = -0.31, 95% HDI ([-0.71, 0.10], (P(θ₁ < 0=0.93). Visually, overlap between the HDI and ROPE suggest uncertainty, however, 42.7% of the posterior distribution exceeded the ROPE falling into practically meaningful regions, suggesting a potential trend in memory probability for the format by age interaction parameter.

##### Memory precision (κ₁)

The posterior mean for memory precision for both the age by time of day (M = 0.07, 95% HDI ([-0.28, 0.39]) (Fig 9C) and age by format (M = 0.08, 95% HDI ([-0.27, 0.43]) (Fig 9D) were centered near 0 and for both interactions and ROPE and HDI intervals overlapped yielding uncertainty. Posterior distributions were split across ROPE thresholds (*p*_meaningful_ = 58.8% and 61.3%, respectively), providing further evidence for neither confirming nor ruling out meaningful interaction effects. Table 2 summarizes ROPE values for the four interaction parameters, as well as decisions based on ROPE and HDI overlap and posterior probabilities.

**Table 2.** Bayesian Parameter Estimates and HDI-ROPE Decisions for Parameter (θ₁, κ₁) Interactions.

| Parameter | Posterior Mean [95% HDI] | ROPE Bounds | $p_{\text{ROPE}}$ | $p_{\text{meaningful}}$ | Decision (ROPE + HDI Overlap) |
| --- | --- | --- | --- | --- | --- |
| **Memory Probability ($\theta_1$)<sup>a</sup> :**<br>**Age × Time of Day** | 0.04 [-0.36, 0.43] | [-0.35, 0.35] | 0.914 | 0.086 | Undecided |
| **Memory Probability ($\theta_1$)<sup>a</sup> :**<br>**Age × Format** | -0.31 [-0.70, 0.10] | [-0.35, 0.35] | 0.573 | 0.427 | Undecided |
| **Memory Precision ($\kappa_1$)<sup>b</sup> :**<br>**Age × Time of Day** | 0.07 [-0.28, 0.39] | [-0.10, 0.10] | 0.412 | 0.588 | Undecided |
| **Memory Precision ($\kappa_1$)<sup>b</sup> :**<br>**Age × Format** | 0.08 [-0.27, 0.43] | [-0.10, 0.10] | 0.387 | 0.613 | Undecided |

HDI = Highest Density Interval; ROPE = Region of Practical Equivalence; $p_{\text{ROPE}}$ = posterior probability that a parameter lies inside the ROPE, $p_{\text{meaningful}}$ = posterior probability that the parameter lies outside the ROPE. Decisions are based on whether the 95% HDI overlaps with the predefined ROPE boundaries (Kruschke, 2018).
<sup>a</sup> Estimates for memory probability ( $\theta_1$ ) are modeled on the logit scale
<sup>b</sup> Estimates for memory precision ( $\kappa_1$ ) are modeled on the log scale

## Discussion

This study used an object-location memory precision task to investigate whether age-related differences in episodic memory precision manifest across contextual factors such as internal biological variables (chronotype) and external task features (e.g., experiment format, time of day). Immediate recall was strongly related to encoding performance, and was relatively consistent across time of day and experimental format. Hierarchical mixture modelling revealed that age-related differences in memory performance were primarily driven by reductions in memory precision rather than the successful retrieval of an item from memory. In contrast, precision and retrieval probability were generally robust to variations in time of day, chronotype alignment, and experiment format. This suggests that participants were able to establish a detailed and precise memory representation of the target objects during encoding which was important for accurate subsequent memory performance largely irrespective of internal or external task features.

The observed age differences in precision were less pronounced than those reported in a number of previous studies, which demonstrated robust age-related declines in the memory precision alongside relatively preserved retrieval success [7,8,42–44]. Methodological differences at the encoding stage between this study and previous precision tasks may have attenuated age effects. Prior studies involved passive viewing of displays with up to three objects presented briefly. In contrast, the learning phase in the present task required more effortful encoding: subjects were shown one object at a time, had to immediately move the object to the studied location, and received feedback. This difference in encoding, particularly the feedback stage, may have reduced the age-related differences in precision at recall that are often found in other object-location memory tasks. The present findings therefore suggest that while memory representations become less precise with age, the magnitude of this decline may depend on task and encoding conditions.

Consistent with prior studies, older adults’ MEQ scores were higher than those of younger adults, indicating a self-reported morning preference among older adults [41,45].Older adults typically report earlier sleep and wake times and tend to perform optimally earlier in the day, whereas younger adults are more likely to exhibit evening preferences [13,14]. As expected, chronotype was associated with the timing of participation: individuals tested in the morning had higher MEQ scores, indicative of a stronger morning preference, whereas those tested in the evening had lower MEQ scores, consistent with a greater evening preference. This pattern suggests that participants’ scheduling choices broadly reflected their underlying circadian preferences. Importantly, there was no significant interaction between age and time of day on MEQ scores, indicating that the relationship between chronotype and testing time was similar in younger and older adults. Thus, although older adults were generally more morning-oriented than younger adults, both age groups showed comparable correspondence between self-reported chronotype and the time at which they elected to complete the task.

Contrary to predictions based on the synchrony effect, completing the task at a time of day that matched participants’ chronotype did not improve memory precision or retrieval probability for either age group. This outcome is supported by overlapping credible intervals and ROPE analysis (91.4% of the posterior distribution within the ROPE), providing strong evidence of a negligible age by time of day interaction. Performance was similar between participants tested at their preferred (matched; N = 134) and non-preferred (mismatched; N = 98) times, contrasting with earlier studies demonstrating synchrony benefits [11,14,41]. Even though participants selected their own testing time, matched and mismatched groups were relatively balanced across age groups, minimising a potential confound between chronotype and testing time. To directly assess synchrony effects, a future experiment could use a within-subjects design that tests participants at their preferred and non-preferred time of day.

Experiment format was associated with lower recall error and higher recognition performance observed in the video (supervised) condition compared to the non-video (unsupervised) condition. However, these behavioural differences were not evident when the data were analyzed using a hierarchical mixture model. Although the model suggested a possible reduction in memory-based responding in the non-video format, the associated uncertainty was substantial, with credible intervals that included zero. For the age and format interaction for memory probability, while the 95% HDI overlapped with the ROPE boundary, posterior probabilities revealed that nearly half of the distribution (*p*_meaningful_ = 42.7%) was in the practically meaningful negative range, suggesting caution in concluding that experimental format has differential effect on older versus younger adults. These findings are broadly consistent with previous studies reporting that supervised and unsupervised online testing generally yield comparable cognitive outcomes, despite occasional differences in overall performance measures [19,20]. Thus, while supervision may confer some practical benefits for data quality, the present findings suggest that estimates of memory precision and retrieval success are relatively robust across testing formats.

The apparent differences observed in behavioural measures (e.g., MAE and recognition performance), should be interpreted with caution because such aggregate measures reflect a combination of underlying cognitive processes, including memory precision, retrieval success, and guessing. In contrast, mixture modelling allows these processes to be estimated separately, providing a more nuanced account of memory performance. Overall, performance remained consistent across participants, regardless of whether the experiment was administered via video or non-video format. The absence of clear format effects on memory precision or memory-based responding therefore suggests that the behavioural differences between video and non-video conditions may not reflect substantive changes in the quality or accessibility of memory representations themselves. Rather, they may arise from broader influences on overall task performance, such as engagement, motivation, environmental distractions, or compliance with task instructions. This interpretation is consistent with previous studies showing that supervised and unsupervised online testing can differ modestly in overall performance measures while nevertheless yielding highly comparable estimates of core cognitive functioning [19,20].

Importantly, this current study extends this literature by including both younger and older adults, and by applying a paradigm that enables dissociation of memory precision from retrieval success. The non-video condition also included periodic attention checks during encoding and recall. Detailed analysis examining the influence of age, time of day, and chronotype on attention-check performance confirmed that participants maintained sufficient attention throughout the task (See Supplementary). Together, these findings suggest that immediate memory retrieval in this paradigm is robust to variations in testing contexts, and that the task can be administered effectively in an online setting, both with and without research supervision.

Unlike the mixture modelling approaches used in many previous studies, the hierarchical modelling implemented here simultaneously estimated individual-and group-level effects while accounting for uncertainty in parameter estimation [32,37]. This was particularly important in the present study because the design incorporated multiple factors, including age group, time of day, experiment format, and chronotype, all of which could potentially contribute to variability in memory performance. Consistent with previous work highlighting substantial heterogeneity in cognitive ageing [40,42], both the primary and secondary models revealed that participants differed substantially in memory precision and retrieval probability. By using the Bayesian HDI-ROPE framework, we were able to evaluate the full posterior distributions of the model parameters, allowing us to separate genuine practical equivalence from potential data noise caused by subject variability. This approach provides a more informative assessment than reliance on conventional null-hypothesis significance testing alone, because it allows evidence for negligible effects to be quantified directly rather than inferred from non-significant results.

Overall, convergence of the hierarchical models and ROPE analyses strengthens the conclusion that memory precision and retrieval probability were largely robust to these contextual factors, even in the presence of marked individual differences.

The ROPEs chosen for these parameters were intentionally conservative, reflecting a preference to avoid overinterpreting weak effects as meaningful. As a consequence, several of the 95% intervals overlapped the ROPE boundaries, leading to categorical classification of “undecided”. To address potential limitations of the HDI-ROPE framework and following recommendations by Etz et al.[46], a sensitivity analysis was performed using 95% quartile intervals (Equal-Tailed Intervals (ETI)) to mitigate possible transformation incoherence. In addition, continuous posterior probabilities associated with these intervals were used to overcome reliance on rigid categorical boundaries. Reassuringly, comparison of the ETI-ROPE and HDI-ROPE approaches revealed no differences in parameter results, indicating that the results were not dependent on the choice of interval estimation method (See Supplementary).

The present study provides a foundation for future research examining how time of day, experiment format, and individual differences in chronotype interact with age under more demanding conditions. In the present study, memory was assessed immediately after learning, placing relatively modest demands on retrieval and minimising forgetting. Previous studies have found that age-related reductions in memory precision become more pronounced as task demands increase, including longer retention intervals and more challenging retrieval conditions [6–8]. Likewise, synchrony effects are most commonly observed in tasks that place greater demands on attention and cognitive control [11,13,14]. Future studies could explore whether the present findings generalise to longer delays, different memory features, and more demanding paradigms, including those involving sleep-dependent consolidation processes [47]. Such studies may help identify the conditions under which ageing, circadian factors, and task context influence both the quality and accessibility of episodic memories.

Although the present findings suggest that chronotype synchrony had little influence on performance, future work could confirm these findings by testing at non-optimal times using a within subject design. Experimentation should include objective physiological measures of arousal (e.g., pupillometry, actigraphy) and circadian markers (e.g., melatonin onset) to confirm chronotype findings based on self reporting. Finally, future memory precision studies that include multiple, interdependent complex factors such as in this study should employ Bayesian hierarchical modeling. Examination of posterior probabilities that represent evidence strength moves beyond categorical null-hypothesis testing. This approach more effectively represents individual variability and group-level effects as well as relationships among variables enabling more precise examination of experimental theories.

## Conclusion

This study demonstrated that age-related differences in episodic memory precision are relatively robust across variations in testing context, including task format, time of day, or an individual’s chronotype. Although older adults exhibited reduced memory precision, there was little evidence that these contextual factors influenced either the quality of memory representations or the probability of successful retrieval. These findings extend previous work demonstrating age-related declines in memory precision [6,7,42] by showing that such effects are detectable under a range of testing conditions, including remote and unsupervised online assessment. In addition, the results suggest that time of testing and chronotype alignment are unlikely to be a major source of variability in immediate object-location memory performance.

Future research should determine whether these findings generalise to more demanding memory conditions, with objective physiological circadian markers (such as pupillometry or melatonin onset) that complement self-reports, explore longer retention intervals (e.g., 24 hours or one week) to examine how these variables impact memory consolidation over time, and evaluate precision tasks with varying characteristics (e.g., perceptual features, cognitive load, speed).

These studies promise further insight into the factors that influence the fidelity and accessibility of memory representations across the adult lifespan.

## Acknowledgements

This research was supported by a Gates Cambridge Trust studentship to JSM and UKRI grant MR/Z505614/1 to JSS. The funders played no role in the study design, data collection and analysis, decision to publish, or preparation of the manuscript. We thank Maheria Abid Rashid for assistance with data collection. For the purpose of open access, the author has applied a Creative Commons Attribution (CC BY) licence to any Author Accepted Manuscript version arising from this submission.

## Supporting information

S1 Fig. Posterior Standard Deviation for Model Parameters across Sample Size. S1 Table. Comparison of HDI and ETI boundaries and ETI-ROPE Decisions for Parameter (θ₁, κ₁) Interactions. *HDI=* High Density Interval, ETI= Equal Tail Interval; ROPE = Region of Practical Equivalence. Decisions are based on whether the 95% ETI overlaps with the predefined ROPE boundaries. *p*_ROPE_ and *p*_meaningful_ are not shown as they are calculated using the entire posterior distribution and therefore remain the same regardless of which interval is used in the ROPE decision.

